# Synthesis and characterization of chloroquine-modified albumin-binding siRNA-lipid conjugates for improved intracellular delivery and gene silencing in cancer cells

**DOI:** 10.1101/2024.10.14.618042

**Authors:** Justin H. Lo, Eva F. Gbur, Nora Francini, Jinqi Ma, Alexander G. Sorets, R. Brock Fletcher, Fang Yu, Richard D’Arcy, Connor G. Oltman, Md. Jashim Uddin, Craig L. Duvall

## Abstract

siRNA therapeutics have considerable potential as molecularly-targeted therapeutics in malignant disease, but identification of effective delivery strategies that mediate rapid intracellular delivery while minimizing toxicity has been challenging. Our group recently developed and optimized an siRNA conjugate platform termed “siRNA-L_2_,” which harnesses non-covalent association with endogenous circulating albumin to extend circulation half-life and achieve tumor-selective delivery without the use of traditional cationic lipids or polymers for transfection. To improve intracellular delivery and particularly the endosomal escape properties of siRNA-L_2_ towards more efficient gene silencing, we report synthesis of siRNA-CQ-L_2_ conjugates, in which chloroquine (CQ), an endosomolytic quinoline alkaloid, is covalently incorporated into the branching lipid tail structure. We accomplished this via synthesis of a novel CQ phosphoramidite, which can be incorporated into a modular siRNA-L_2_ backbone using on-column solid-phase synthesis through use of asymmetric branchers with levulinyl-protected hydroxide groups that allow covalent addition of pendant CQ repeats. We demonstrate that siRNA-CQ-L_2_ maintains the ability to non-covalently bind albumin, and with multiple copies of CQ, siRNA-CQ-L_2_ mediates higher endosomal disruption, cellular uptake/retention, and reporter gene knockdown in cancer cells. Further, in mice, the addition of CQ did not significantly affect circulation kinetics nor organ biodistribution and did not produce hematologic or organ-level toxicity. Thus, controlled, multivalent conjugation of albumin-binding siRNA-L_2_ to endosomolytic small molecule compounds holds promise in improving siRNA-L_2_ knockdown potency while maintaining albumin-binding properties and overall safety.

## 1. Introduction

RNA interference mediated by small interfering RNA (siRNA) holds great promise as molecularly-targeted therapy in cancer.^1^ Numerous delivery systems have been developed to convey siRNA payloads to tumor cells, though many rely upon cationic lipids or polymers that can lead to systemic toxicity or inflammation,^2,3^ and none have yet been translated into clinical practice for oncology. As an alternative, direct siRNA conjugates are being developed in which the siRNA molecule itself is elaborated with covalent modifications that mediate delivery, enabled by crucial siRNA-stabilizing modifications that prevent nuclease degradation of unshielded siRNA bases.^4^ Indeed, five of the six FDA-approved siRNA therapeutics are GalNAc-conjugated siRNA molecules that target hepatocytes.^5^ Unlike traditional cationic delivery vehicles, siRNA conjugates typically lack dedicated endosome escape functionality and consequently tend to accumulate in endosomal “depots,” leading to prolonged slow release into the cytoplasm that can be favorable for the treatment of benign conditions.^6^ However, development of siRNA conjugates for cancer applications necessitates rapid endosomal escape, since the dilutional effect of rapid cell division in cancer cells is the primary factor limiting the activity of siRNA therapeutics.^7^ There have been efforts to overcome endosomal barriers to siRNA conjugate delivery in extrahepatic tissues, for example by covalent conjugation of ionizable lipids, borrowing from the lipid nanoparticle repertoire.^8^ Unfortunately, this approach led to toxicity *in vivo*, evidenced by non-specific gene expression modulation, leaving an unmet need for methodologies for improving cytoplasmic delivery of siRNA conjugates without significantly limiting the therapeutic index as with traditional cationic delivery vehicles.

To develop siRNA conjugates for cancer applications, our group has modified siRNA with twin lipid-terminated tails (siRNA-L_2_) that non-covalently bind to endogenous albumin.^9,10^ Optimization of this “albumin hitchhiking” approach results in extended circulation kinetics, tumor tropism, and gene silencing in mouse models of triple negative breast cancer.^10^ We considered that the existing siRNA-EG18-L_2_ structure may achieve more efficient endosomal escape and thus greater potency through incorporation of a dedicated endosomal escape mechanism.

One method of interest for enhancing endosomal escape of nucleic acid therapeutics is to co-administer chloroquine (CQ), a quinoline alkaloid originally developed and used clinically as an antimalarial agent. CQ disrupts endosomal acidification as a weak base and directly interacts with the endosomal membrane to promote permeabilization.^11^ Co-administration of free chloroquine enhances endosomal disruption and reporter gene silencing of cholesterol-conjugated siRNA.^12^ However, this approach has not seen clinical translation given the excessive doses of co-administered free CQ required, which has a narrow therapeutic window, does not target to sites of disease, and does not match the pharmacokinetics of siRNA therapeutics. We hypothesized that covalent addition of chloroquine to the siRNA-L_2_ conjugate backbone would co-localize its effects to the cellular endosomes containing siRNA-L_2_, improving the endosomal escape properties and potency of siRNA-L_2_ while maintaining its essential albumin-binding functionality and minimizing non-specific toxicity.

## 2. Materials and Methods

### Desulfation of Hydroxychloroquine

Hydroxychloroquine (HCQ) sulfate (TCI) was desulfated according to a method adapted from Yu et al.^13^ Briefly, 6.0 g of HCQ sulfate was dissolved in 40 mL of water. 5.0 mL of NH_4_OH was added dropwise while stirring vigorously. After 30 minutes, 40 mL of dichloromethane was added. After 10 more minutes, the solution was transferred to a separatory funnel, and the organic layer was collected and washed with brine, then dried with sodium sulfate. Dichloromethane was evaporated *in vacuo*, yielding HCQ.

### Synthesis of Chloroquine Phosphoramidite

2 mmol of desulfated HCQ was dissolved in 3 mL of anhydrous dichloromethane. 2.8 mmol of N,N-diisopropylethylamine (DIPEA, Millipore Sigma) was added while stirring at room temperature. 1.8 mmol of 2-cyanoethyl N,N-diisopropylchlorophosphoramidite (Millipore Sigma) was added dropwise and the reaction was stirred at room temperature for 1 hour under nitrogen. The reaction mixture was extracted against brine twice and then dried with sodium sulfate for 30 minutes followed by evaporation of dichloromethane *in vacuo*. The product was resuspended in 3:1 anhydrous dichloromethane:acetonitrile and transferred into an oven-dried glass vial for use on a Mermade oligonucleotide synthesizer.

### NMR Confirmation of Chloroquine Phosphoramidite

^1^H-NMR (400 MHz, DCM-*d*_*2*_) 1.00 (s, 3H), 1.02 (s, 3H), 1.04 (s, 3H), 1.17 (s, 3H), 1.19 (s, 3H), 1.20 (s, 3H), 1.60–1.63 (m, 2H), 1.65–1.69 (m, 2H), 2.52–2.71 (m, 11H), 3.60-3.64 (m, 2H), 3.74-3.80 (m, 3H), 6.48 (d, *J* = 8.5 Hz, 1H), 7.43 (dd, *J* = 9.0, 2.2 Hz, 1H), 7.82 (dd, *J* = 8.9, 2.1 Hz, 1H), 7.92 (d, *J* = 2.5 Hz, 1H), 8.51 (d, *J* = 8.4 Hz, 1H). ^31^P-NMR (400 MHz, DCM-*d*_*2*_) 147.57 (s, 1P).

### Synthesis of Chloroquine-modified siRNA-L_2_

The on-column synthesis of the siRNA-L_2_ backbone on a Mermade oligonucleotide synthesizer has been described in Hoogenboezem et al.^10^ Briefly, the sense strand of the siRNA sequence is synthesized using Zipper modifications on the RNA bases (alternating 2’ O-methyl and 2’-fluoro modifications), along with phosphorothioate linkages between the three terminal bases on both the 5’ and 3’ ends of both strands. The EG18-L_2_ tail, consisting of a symmetrical brancher (ChemGenes) followed by 3 repeats of (ethylene glycol [EG])_6_ and a terminal stearic acid (C_18_) on each branch, is appended to the 5’ end of the sense (passenger) siRNA strand, with each group linked by a phosphorothioate bond for stability. In contrast to the previously published structure, we have interpolated the addition of 1-3 instances of an asymmetric brancher (ChemGenes CLP-7169), in which one arm bears a conventional trityl protecting group while the other is protected with a levulinyl group (**Figure 1B**). Initially, the trityl arm is extended with the remainder of the L_2_ tail (symmetric brancher followed by 3 instances of ethylene glycol (EG)_6_ and terminal stearyl groups, all linked by phosphorothioate bonds. Then, we deprotected the pendant levulinyl-protected hydroxyl group with hydrazine hydrate (Millipore Sigma) in 1:1 acetic acid:pyridine, washed with 1:1 acetic acid:pyridine followed by anhydrous acetonitrile x4, and finally conjugated the chloroquine phosphoramidite. siRNA-CQ-L_2_ conjugates were cleaved from the columns using 1:1 ammonium hydroxide:methylamine (40% in H_2_O, Millipore Sigma) followed by desalting, HPLC purification (Shimadzu), and removal of organic and aqueous solvents using a Thermo SpeedVac SPD120, as previously described. siRNA-CQ-L_2_ was resuspended in sterile saline and molecular weight was confirmed using liquid chromatography-mass spectrometry (LCMS) as previously described.

**Figure 1.**
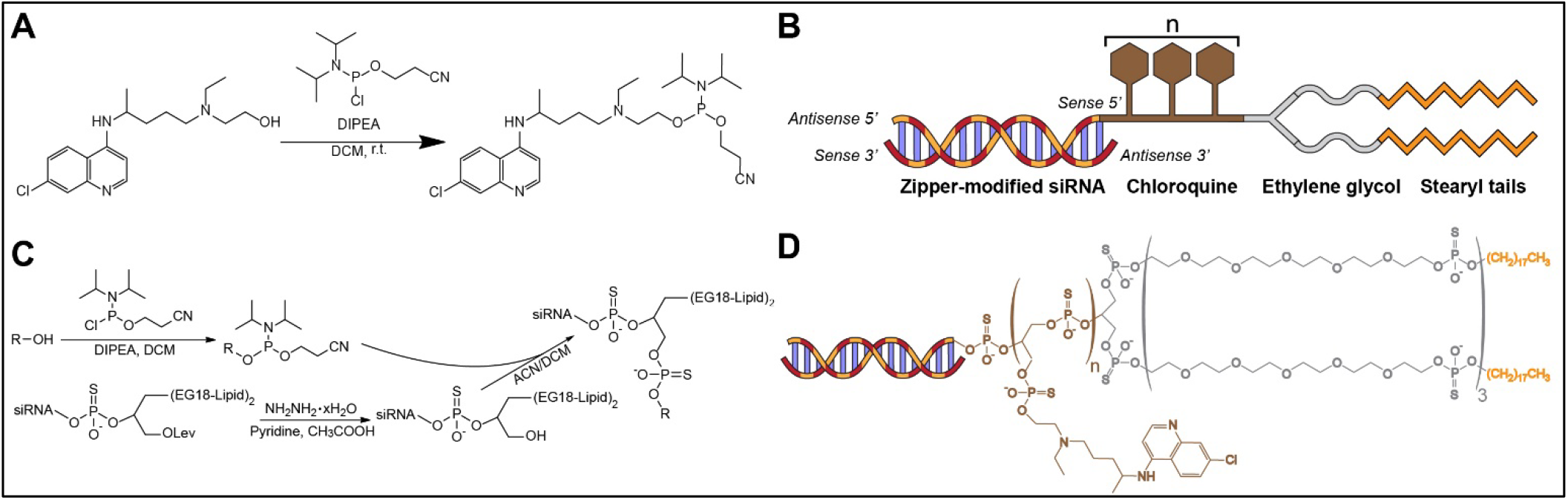
Synthesis of chloroquine phosphoramidite and chloroquine-modified siRNA-lipid conjugates. **(A)** Synthesis scheme for CQ phosphoramidite. **(B)** Schematic of siRNA-CQ-L_2_ structure. **(C)** Summary of on-column solid-phase synthesis of siRNA-CQ-L_2_. **(D)** Chemical structure of siRNA-CQ-L_2_.

### Molecular weight and AlogP calculations

Molecular weights were calculated using ChemDraw v22.2 (Revvity) and AlogP was calculated using BIOVIA Draw 2022 (Dassault Systèmes).

### Dynamic Light Scattering

Solutions of the specified siRNA-L_2_ formulations were diluted to 2 μM in DPBS, pH 7.4 (Gibco) and mixed with a final concentration of 2 μM human serum albumin (Millipore Sigma). Hydrodynamic diameter was measured in a ZEN2112 quartz cuvette using a Malvern ZS90 dynamic light scattering instrument and ZS Xplorer software (Malvern Pananalytical).

### Biolayer Interferometry

Binding kinetics of siRNA-CQ-L_2_ to human serum albumin were measured by biolayer interferometry (BLI) at 30 °C, 1000 rpm using an Octet R8 (Sartorius). Biotinylated human serum albumin (25 μg/mL) was loaded for 600 sec on Octet SA Biosensors (Sartorius). Biosensor association to siRNA-L_2_ or siRNA-CQ-L_2_ duplexes diluted to the specified concentrations (1 to 0.00625 μM) in DPBS was measured for 300 sec, followed by 180 sec of baseline and 300 sec of dissociation measurements. The binding values were measured using Octet Data Analysis HT Software. Interstep correction was performed by aligning to the dissociation step, and noise filtering was performed. Global analysis was performed to derive constants simultaneously from all tested analyte concentrations.

### Critical Micelle Concentration

In a black 384-well plate (Greiner), a 1:2 serial dilution of the specified siRNA-L_2_ conjugates was made in 50 μL of DPBS without calcium or magnesium, pH 7.4 (Gibco), spanning from 20 to 0.01 μM. To each well, 5 μL of 0.1 mg/mL Nile Red in acetone (TCI) was added. The plate was covered in foil and agitated at 37 °C for 2 hours to evaporate the acetone. Fluorescence was quantified on a Tecan Infinite M1000 Pro plate reader using excitation 535 ± 10 nm and emission 610 ± 10 nm. Logarithmic regression for the lower and upper data points was performed using Microsoft Excel, and the intersection of these curve fits was taken to be the critical micelle concentration (CMC).

### Gal8 Endosomal Escape Assay

Generation of MDA-MB-231 cells expressing the Gal8-YFP fusion protein has been previously described in Kilchrist et al.^14^ A black 96-well plate was coated with 50 μg/mL rat tail collagen (Gibco) in 20 mM sterile acetic acid for 30 min at 37 °C and subsequently rinsed with sterile DPBS (Gibco). Then, Gal8-YFP MDA-MB-231 cells were plated at 10,000 cells per well in high-glucose DMEM (Gibco) supplemented with 10% fetal bovine serum (FBS, Avantor) and penicillin/streptomycin (Gibco) and allowed to attach for 24 hours. Wells were rinsed with 100 μL of Opti-MEM (Gibco) and then treated with (1) media only, (2) lipofectamine 2000 (Invitrogen) with zipper-modified luciferase siRNA (100 nM, synthesized on Mermade) according to manufacturer instructions, (3) luciferase siRNA-EG18-L_2_ at 1 μM, (4) the same supplemented with 3 μM free chloroquine diphosphate (Millipore Sigma), (5) luciferase siRNA-CQ_1_-EG18-L_2_ at 1 μM, (6) luciferase siRNA-CQ_2_-EG18-L_2_ at 1 μM, or (7) luciferase siRNA-CQ_3_-EG18-L_2_ at 1 μM, n=4 per condition. After incubation for 24 hours, wells were supplemented with NucBlue live cell stain (Invitrogen) in Opti-MEM according to manufacturer instructions and imaged on a Nikon Eclipse Ti confocal microscope using Nikon Elements software v. 4.01. Timelapse imaging was performed using an on-stage plate warmer supplied with humidified 5% CO_2_ in room air to maintain physiologic conditions. In this experiment, Gal8 MDA-MB-231 cells were treated with Cy5-labeled siRNA-CQ_1_-L_2_ (1 μM) or lipofectamine RNAiMax (Invitrogen) with zipper-modified siRNA (100 nM). In all cases, images were analyzed using MATLAB R2023a (Mathworks) using an image analysis algorithm as previously described by our lab.^15^

### *In vitro* toxicity assessment

MDA-MB-231 cells were plated at 5,000 cells per well in a black collagen-coated plate (see above) and incubated for 24 hours prior to treatment with the indicated siRNA-L_2_ formulations at 250, 500, or 1000 nM for 24 hours in Opti-MEM (n=5 per condition). Cell viability was assessed using the CellTiter Glo assay (Promega) according to manufacturer instructions and luminescence was quantified on a Tecan Infinite M1000 Pro plate reader. Relative luminescence was normalized to media-only controls.

### Flow Cytometry for Uptake

MDA-MB-231 cells expressing GFP were plated at 5,000 cells per well in a black 96-well collagen-coated plate and incubated for 24 hours prior to treatment with the indicated Cy5-labeled siRNA-L_2_ formulations at 100 nM for either 1 or 4 hours in Opti-MEM. Cells were then trypsinized and resuspended as a single-cell suspension in 200 μL of PBS + 2% FBS in a 96-well U-bottom plate. Flow cytometric analysis was performed on a Guava easyCyte HT flow cytometer (Cytek). Cells were gated based on FSC/SSC and subsequently for GFP positivity. The geometric mean for Cy5 fluorescence was computed using FlowJo v. 10 software.

### Reporter Gene Silencing

MDA-MB-231 cells expressing firefly luciferase were plated at 5,000 cells per well in a black 96-well collagen-coated plate and incubated for 24 hours prior to treatment with the indicated siRNA-L_2_ formulations at 250, 500, or 1000 nM for 24 hours in Opti-MEM (n=5 per condition). 150 μg/mL D-Luciferin (Pierce) was added to the wells and luminescence was quantified on a Tecan Infinite M1000 Pro plate reader. Relative luminescence was normalized to media-only controls.

### Circulation Kinetics (Peptide Nucleic Acid Hybridization Assay)

Female NCr/nu mice (Envigo) were injected with siRNA-L_2_ or siRNA-CQ-L_2_ at 1 mg/kg body weight (n=4 each). 10 μL of blood was collected into heparinized capillaries by tail vein nick using a 30-gauge needle at 1 minute, 30 minutes, 1 hour, 3 hours, 6 hours, and 24 hours. The peptide nucleic acid hybridization assay was performed in a method adapted from Godinho et al.^16^ Cy3-labeled peptide nucleic acid probes complementary to the luciferase antisense siRNA strand were purchased from PNA Bio. 5 μL of whole blood from each sample was homogenized in 300 μL homogenization buffer (QuantiGene Homogenizing Solution, Thermo Fisher) with 0.5 mg/mL Proteinase K. Standards containing 1:2 serial dilutions from 10,000 to 156 fmol of siRNA were diluted into untreated whole blood processed in the same way. Samples were incubated at 65 °C for 1 hour with periodic vortexing every 15 min. Samples were then centrifuged at 15,000 ×g for 15 min. 200 μL of each sample was carried forward and SDS was precipitated through addition of 20 μL 3M KCl. Samples were centrifuged at 4000 ×g for 15 min, transferred to a new tube, and centrifuged again at 4000 ×g for 5 min. 150 μL of supernatant was then mixed with 100 μL of 50 mM Tris with 10% acetonitrile, pH 8.8 and 2 μL of Cy3-PNA probe at 5 μM. Samples were then incubated at 90 °C for 15 min followed by 50 °C for 15 min. Samples were mixed thoroughly and 150 μL was transferred into a vial for HPLC analysis. HPLC analysis was performed on a Shimadzu X using a DNAPac PA100 anion exchange column (Thermo Fisher). The mobile phase buffers were buffer A (50% water, 50% acetonitrile, 25□mM Tris-HCl pH 8.5, and 1□mM EDTA) and buffer B (800□mM NaClO_4_ in buffer A). Concentrations were determined through measurement of the area under the curve of the fluorescent peak (excitation 550 nm, emission 570 nm) at ∼6 minutes corresponding to hybridized probe and antisense strand and converted to absolute units using the standard curve samples.

### Organ Biodistribution

Wild-type FVB mice were injected with saline control, siRNA-L_2_, siRNA-CQ_2_-L_2_, or siRNA-CQ_3_-L_2_ at 5 mg/kg body weight (both males and females included in each treatment group). After 48 hours, mice were euthanized and blood was collected via cardiac puncture. Whole blood samples were collected in potassium EDTA-coated (lavender top) tubes for complete blood count analysis, while serum samples were prepared by allowing blood to clot in an uncoated tube at room temperature for 1 hour followed by centrifugation at 10,000 ×g for 10 mins x2. Blood counts were performed by the Vanderbilt TPSR Core while serum chemistries were performed by Antech GLP. The specified organs were taken during necropsy and fixed in 10% neutral buffered formalin for 72 hours, embedded in paraffin, sectioned, and stained with hematoxylin and eosin using standard procedures. Slides were scanned using a Leica SCN400 slide scanner by the Vanderbilt Digital Histology Shared Resource. Tissue accumulation of siRNA treatments was performed using the PNA hybridization assay, with ∼5-10 mg of each organ homogenized into 300 μL homogenization buffer using stainless steel beads (Qiagen) in a TissueLyser II tissue homogenizer (Qiagen) for 3 minutes and then proceeding identically to the PNA hybridization assay protocol above.

### Statistical Analysis

Data were analyzed using the specified statistical tests in GraphPad Prism 10 (GraphPad/Dotmatics).

All animal experiments were performed in accordance with the Vanderbilt Animal Care and Use Program.

## 3. Results

### 3.1 Synthesis of chloroquine phosphoramidite and chloroquine-modified siRNA-lipid conjugates

In order to incorporate chloroquine directly into the backbone of the siRNA-L_2_ conjugate using an oligonucleotide synthesizer, we devised a method for synthesizing a chloroquine phosphoramidite. This two-step procedure consists of first desulfating hydroxychloroquine sulfate followed by reaction of hydroxychloroquine with 2-cyanoethyl N,N-diisopropylchlorophosphoramidite (**Fig. 1A**; see Methods for more details). We confirmed successful synthesis of this novel phosphoramidite through electrospray ionization (ESI) mass spectrometry (**Fig. S1**) and ^1^H NMR (see description in Materials and Methods).

Next, we sought to use this phosphoramidite to incorporate chloroquine into the siRNA-L_2_ conjugate backbone. Recent optimization of the branching structure and length of the ethylene glycol^17^ spacer preceding the L_2_ lipid tails led to identification of a lead formulation termed siRNA-EG18-L_2_, and hereafter, siRNA-L_2_ will be used to refer specifically to this EG18 formulation.^10^ The siRNA-L_2_ conjugate structure on the 5’ end of the siRNA sense strand features a symmetric brancher followed by three consecutive ethylene glycol_6_ (EG6) spacers and twin stearic acid tails (L_2_), all joined by phosphorothioate linkages (**Fig. 1B**). To maintain the ability of this tail structure to engage with albumin, while also avoiding modification of either the 3’ end of the sense strand (which is in steric proximity of the critical 5’ phosphate on the antisense strand) or the antisense strand itself, we elected to integrate the chloroquine molecules between the RNA bases and the EG18-L_2_ tails on the 5’ end of the sense strand (**Fig. 1B**). To create this design, we performed on-column solid-phase synthesis of the complete siRNA-L_2_ structure with additional asymmetric brancher units (n=1, 2, or 3) between the siRNA bases and the symmetric branching tail structure. Subsequent deprotection of the asymmetric levulinyl-protected hydroxyl groups allowed conjugation of pendant chloroquine groups using our chloroquine phosphoramidite (**Fig. 1C**), resulting in the final chemical structure depicted in **Fig. 1D**. The final product, termed siRNA-CQ_n_-L_2_ where n=1, 2, or 3, was purified using HPLC, and the correct molecular weight was confirmed using liquid chromatography-mass spectrometry (LC-MS, **Fig. S2**).

### 3.2 Physical properties of chloroquine-modified siRNA-lipid conjugates

We next sought to characterize the physical properties of the siRNA-CQ-L_2_ conjugates; these are summarized in **Table 1**. Given the hydrophobicity of chloroquine, we calculated the AlogP values for the extended tail structure and the overall sense strand, showing that while the tail is predicted to be more hydrophobic, as anticipated, the overall siRNA sense strand remains hydrophilic based on the overall AlogP values, in agreement with our observation that the siRNA-CQ-L_2_ single strands and annealed duplexes are readily soluble in aqueous solution. Accordingly, the critical micelle concentration (CMC) was essentially unchanged from that of the parent siRNA-L_2_ conjugates regardless of the number of chloroquine molecules added, in the range of 3-5 μM (**Table 1, Fig. 2A**). We did not observe aggregates in the complexes formed between human serum albumin (HSA) and siRNA-CQ-L_2_ at 2 μM, though there was a small increase in the hydrodynamic diameter as measured by dynamic light scattering (DLS) indicative of nanocomplex formation between the albumin and siRNA. Finally, we measured the binding affinity of siRNA-CQ-L_2_ for HSA using biolayer interferometry (BLI, **Fig. 2B**). We noted a similar but slightly higher K_D_ in the siRNA-CQ-L_2_ formulations compared to parent siRNA-L_2_. The association constant was unchanged among all formulations, and this small difference in K_D_ instead reflected an increase in the dissociation constant (k_d_).

**Table 1.** Summary of physical properties of chloroquine-modified siRNA-lipid conjugates. For properties marked with an asterisk (*), the sense strand for luciferase siRNA is included in the analysis.

|  | siRNA-EG18-L<sub>2</sub> | siRNA-CQ<sub>1</sub>-EG18-L<sub>2</sub> | siRNA-CQ<sub>2</sub>-EG18-L<sub>2</sub> | siRNA-CQ<sub>3</sub>-EG18-L<sub>2</sub> |
| --- | --- | --- | --- | --- |
| M.W. (tail only), g/mol | 3028 | 3610 | 4193 | 4776 |
| M.W. (with siRNA*), g/mol | 9292 | 9875 | 10458 | 11040 |
| AlogP (tail only) | 8.33 | 11.42 | 14.52 | 17.61 |
| AlogP (with siRNA*) | -10.40 | -7.31 | -4.21 | -1.12 |
| Hydrodynamic diameter of HSA:siRNA, nm | 9.67 | 10.27 | 10.99 | 11.88 |
| CMC, μM | 4.2 | 3.4 | 3.5 | 4.1 |
| Albumin K<sub>D</sub>, nM | 2.3 ± 0.1 | 6.0 ± 0.1 | 18.2 ± 0.1 | 5.9 ± 0.2 |
| Albumin k<sub>a</sub>, M<sup>-1</sup>s<sup>-1</sup> | 3.1 ± 0.0 $\times 10^4$ | 2.5 ± 0.0 $\times 10^4$ | 3.0 ± 0.0 $\times 10^4$ | 3.3 ± 0.0 $\times 10^4$ |
| Albumin k<sub>d</sub>, s<sup>-1</sup> | 7.0 ± 0.4 $\times 10^{-5}$ | 14.9 ± 0.4 $\times 10^{-5}$ | 54.5 ± 0.3 $\times 10^{-5}$ | 19.9 ± 0.3 $\times 10^{-5}$ |

**Figure 2.**
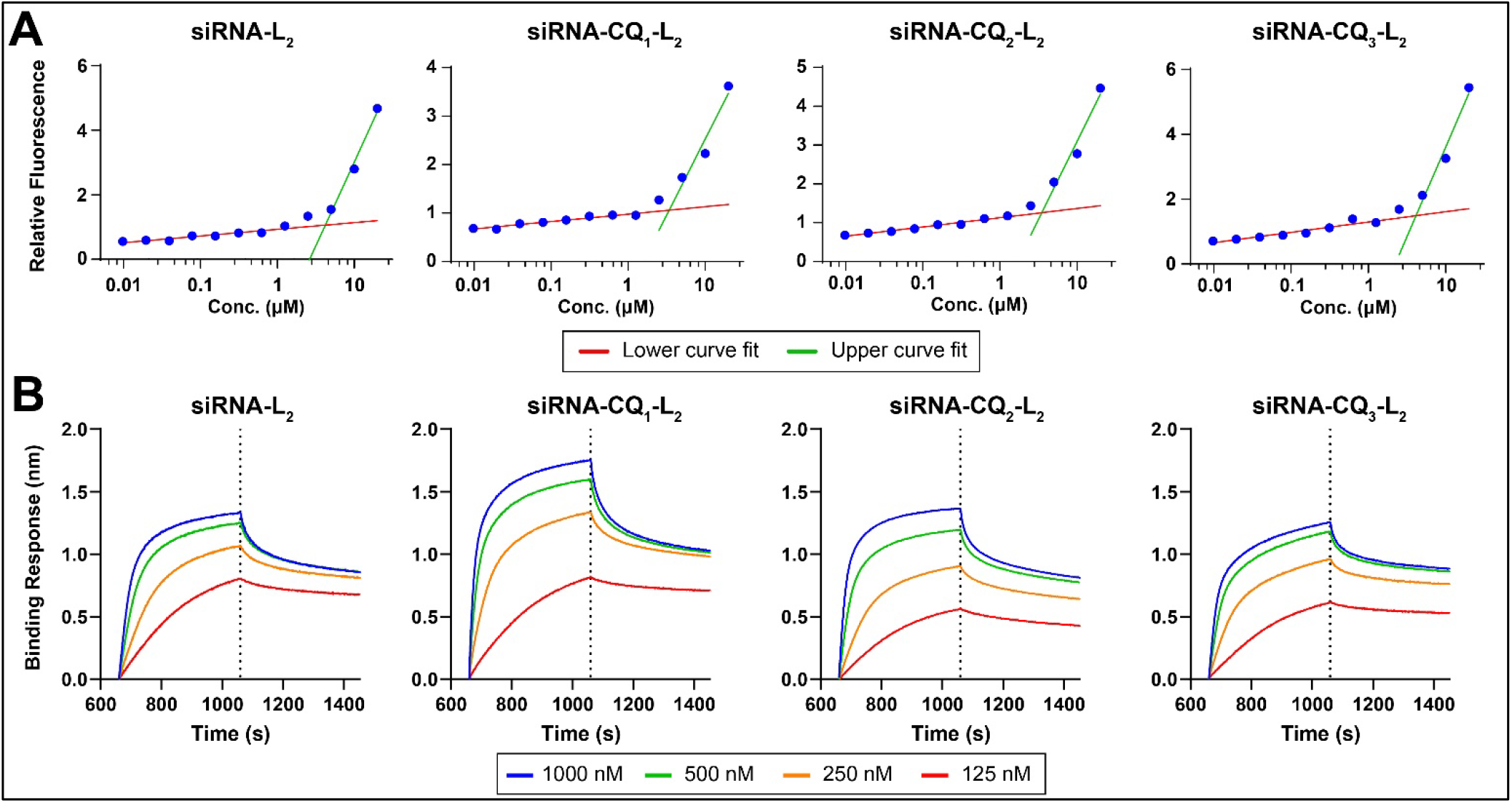
Determination of critical micelle concentration and albumin-binding affinity of siRNA-CQ_n_-L_2_. **(A)** Fluorescence of Nile Red incubated with varying concentrations of siRNA-L_2_ or siRNA-CQ_n_-L_2_. Datapoints represent average of two technical replicates. Red and green curves represent logarithmic regression fits for the 0.01-1.25 μM ranges and 5-20 μM ranges, respectively. The intersection of these curves was taken to be the inflection point representing the CMC. **(B)** Biolayer interferometry curves for binding of varying concentrations of siRNA-L_2_ or siRNA-CQ_n_-L_2_ to immobilized biotinylated human serum albumin, showing association (left) and dissociation (right) phases, separated by the dotted line.

### 3.3 Chloroquine-modified siRNA-lipid conjugates mediate active endosomal escape

The parent siRNA-L_2_ molecule, when administered without use of a transfection reagent, tends to accumulate in endolysosomal compartments, as visualized at 24 hours using co-localization of Cy5-labeled siRNA-L_2_ and Lysotracker dye (**Fig. 3A, left**). Co-administration of free chloroquine at 50 μM results in disruption of normal endosomal maturation and earlier cytoplasmic accumulation of siRNA-L_2_ **(Fig. 3A, right**).

**Figure 3:**
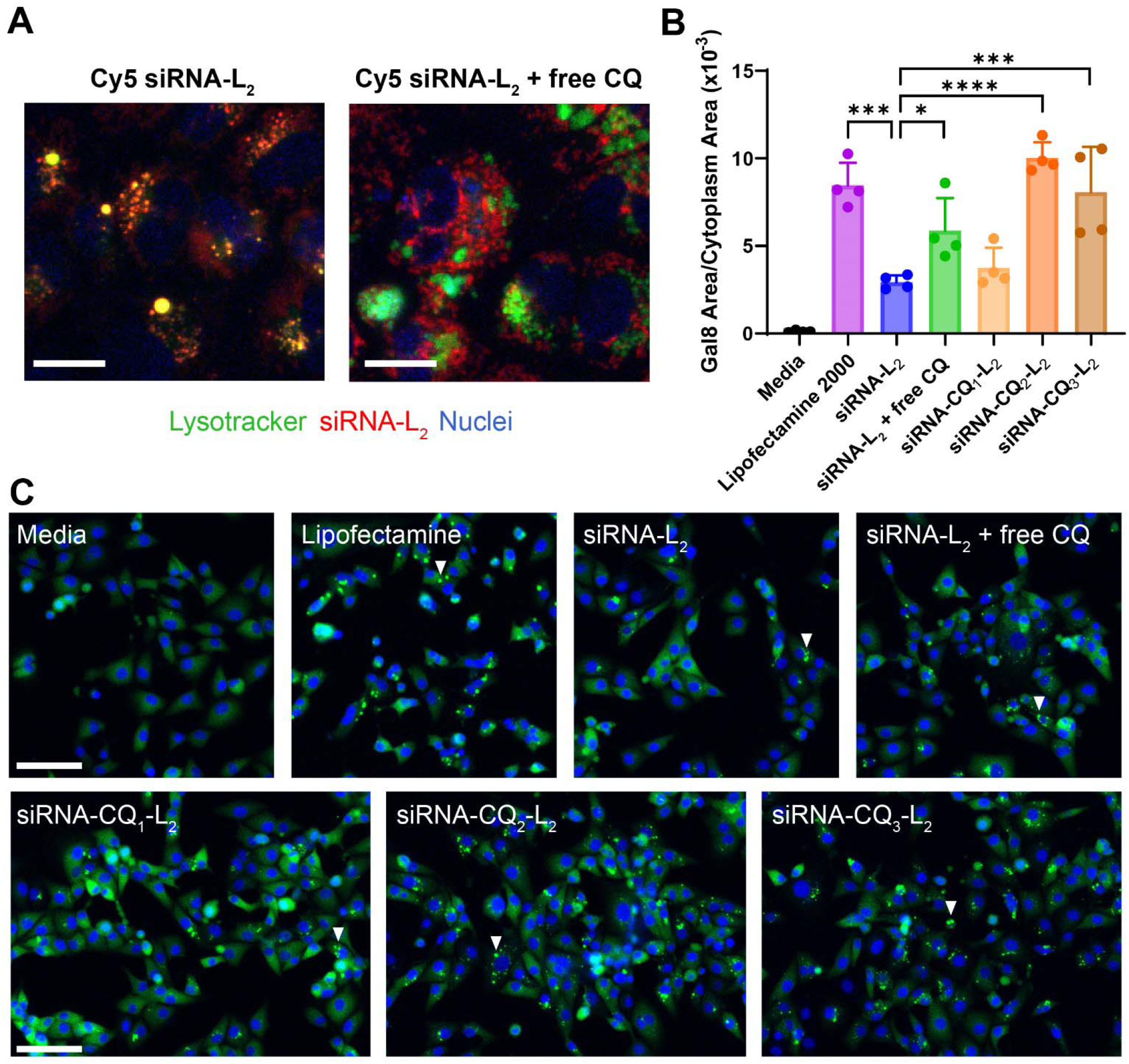
Endosomal escape properties of free chloroquine and siRNA-CQ_n_-L_2_. **(A)** Effect of free CQ on siRNA-L_2_ release into cytoplasm in HuCC-T1 cells *in vitro*. Scale bar = 20 μm. **(B)** Quantification of area of Gal8 foci in Gal8-YFP MDA-MB-231 cells incubated with media only, lipofectamine RNAiMax (positive control), siRNA-L_2_ alone, siRNA-L_2_ with free 3 μM CQ, or siRNA-CQ_n_-L_2_. Asterisks indicate statistical significance by one-way ANOVA: *: p<0.05, ***: p<0.001; ****: p<0.0001. **(C)** Representative confocal microscopy of Gal8-YFP MDA-MB-231 cells incubated with the specified conditions. White arrowheads indicate examples of Gal8 foci, in contrast to diffuse cytoplasmic fluorescence, which is only used to quantify cytoplasmic area. Scale bars = 50 μm.

We directly visualized and quantified the level of endosomal disruption mediated by siRNA-CQ-L_2_ versus the parent siRNA-L_2_ conjugate using the Gal8-YFP assay developed previously in the Duvall lab, in which endosomal disruption leads to localization of fluorescently-tagged Gal8 into bright foci that can be quantified objectively through image processing (**Fig. 3B-C**).^14,15^ In Gal8-YFP MDA-MB-231 cells, siRNA-L_2_ produced low levels of Gal8 foci while addition of free CQ or incubation with siRNA-CQ_2_-L_2_ or siRNA-CQ_3_-L_2_ led to statistically significantly increased Gal8 signal, quantified as total area of Gal8 foci normalized to the total cytoplasmic area. In contrast to lipofectamine RNAiMax transfection reagent, which produces robust Gal8 foci within a few hours and toxicity if left on cells for an extended period of time (over 4-6 hours), siRNA-CQ-L_2_ produces foci more gradually (**Fig. S3**).

### 3.4 Chloroquine-modified siRNA-L_2_ conjugates exhibit improved uptake patterns and gene silencing profiles without increasing toxicity *in vitro*

Next, we determined the effects of covalent addition of chloroquine on siRNA-L_2_ function *in vitro*. First, we quantified uptake of siRNA-CQ-L_2_ as compared to parent siRNA-L_2_ or siRNA-L_2_ with co-administered free chloroquine (50 μM) at a fixed 100 nM dose of siRNA. For these studies, siRNA-L_2_ or siRNA-CQ_1-3_-L_2_ sense strands were annealed to 5’ Cy5-labeled antisense strand, and equal fluorescence of each resulting double-stranded siRNA conjugate was confirmed. We treated adherent MDA-MB-231 triple negative breast cancer cells for 1 hour (**Fig. 4A-B**) or 4 hours (**Fig. 4C-D**) at 37 °C. At both timepoints, we observed significant increases in uptake with the siRNA-CQ_2_-L_2_ and siRNA-CQ_3_-L_2_ formulations, while free chloroquine and CQ_1_-L_2_ only led to increased uptake at the earlier timepoint. We also performed the same treatments for 2 hours at 4 °C to isolate increased surface binding from active internalization or retention; no increases in fluorescent uptake were seen for any of the siRNA-CQ-L_2_ conditions in this setting (**Fig. S4**).

**Figure 4:**
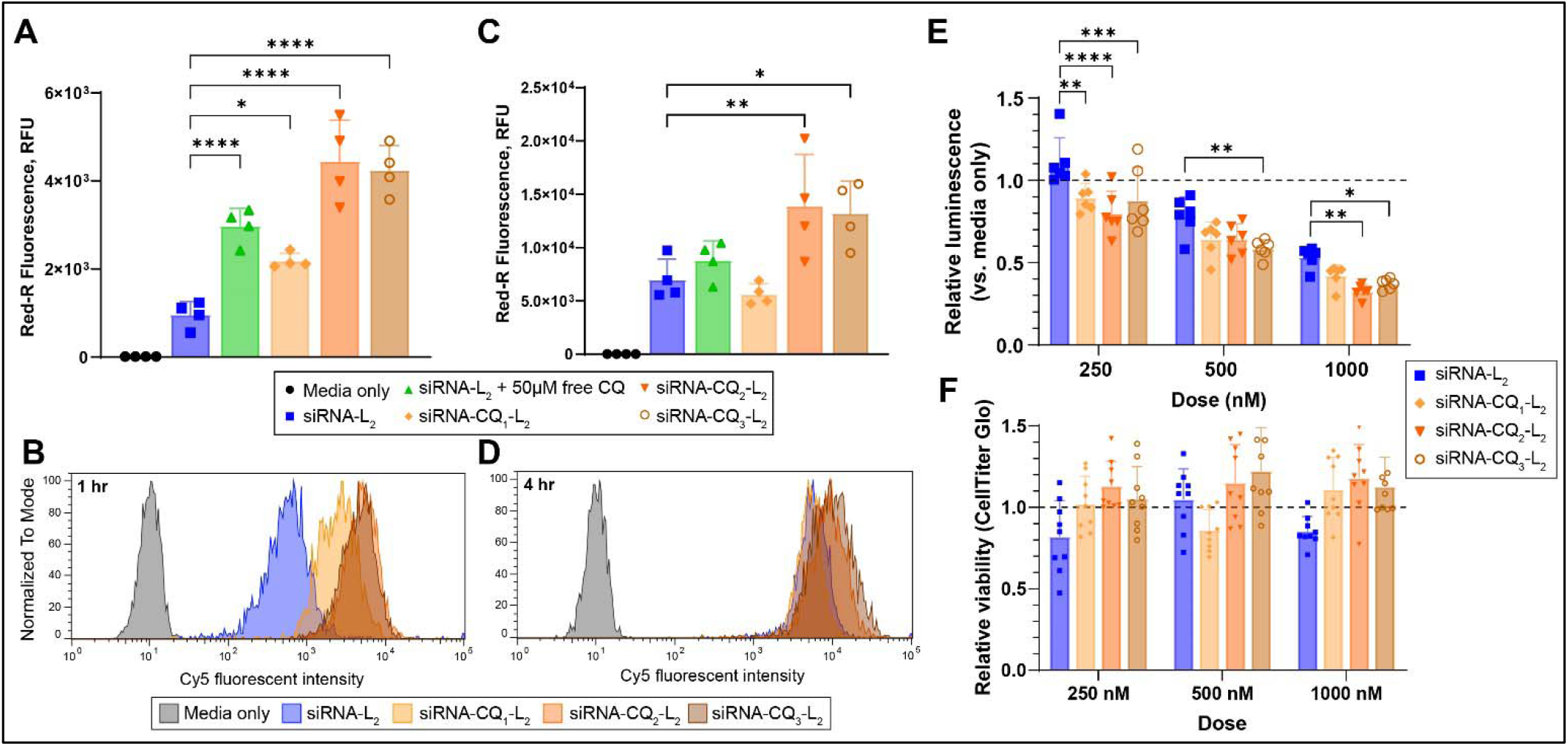
*In vitro* characterization of siRNA-CQ-L_2_ uptake, gene silencing, and toxicity. **(A-D)** Flow cytometry analysis of uptake of Cy5-labeled siRNA-L_2_ (without or with free chloroquine) or Cy5-labeled siRNA-CQ_n_-L_2_ in MDA-MB-231 cells. Geometric means of Cy5 fluorescence (n=4 per condition) and representative Cy5 fluorescence histograms are shown following 1 hour **(A-B)** or 4 hours **(C-D)** of incubation with 100 nM of siRNA conjugates or media only at 37 °C. Asterisks indicate statistical significance by one-way ANOVA with Dunnett’s post-test comparing experimental groups against siRNA-L_2_. *: p<0.05, **: p<0.01; ****: p<0.0001. **(E)** Firefly luciferase silencing by luciferase mRNA-targeting siRNA-L_2_ or siRNA-CQ_n_-L_2_ following 24 hour incubation of Luc-expressing MDA-MB-231 cells at the specified doses. *: p<0.05, **: p<0.01; ***: p<0.001; ****: p<0.0001 **(F)** Cell viability measured by CellTiter Glo following 24 hour incubation of non-Luc-expressing MDA-MB-231 cells (to avoid conflict with the luminescent viability assay) with siRNA-L_2_ or siRNA-CQ_n_-L_2_ at the specified doses. Asterisks indicate statistical significance by two-way ANOVA with Dunnett’s post-test comparing experimental groups against siRNA-L_2_. **: p<0.01.

Next, we sought to determine whether these CQ-driven changes in endosome disruption and intracellular retention impact gene silencing potency and efficacy. We have previously observed that optimal knockdown with siRNA-L_2_ in the absence of any transfection agents (“carrier-free” delivery) is achieved with 1000 nM doses after 72 hours when treating adherent cells. We hypothesized that the extended time to onset and relatively high dose requirements reflects dose sequestration and slow release form endosomes. Therefore, we compared silencing of reporter firefly luciferase in adherent MDA-MB-231 cells at varying doses in the range of 250-1000 nM at a much earlier 24 hour timepoint, with luminescence normalized to media-only controls. At every dose level, there was a trend towards greater knockdown with increasing chloroquine content, with siRNA-CQ_3_-L_2_ consistently showing statistically significant improvements in luciferase silencing compared to siRNA-L_2_ at all doses tested (**Fig. 4E**). This effect was not due to non-specific cell toxicity, as viability of cells treated with any of these conditions was comparable as measured by CellTiter Glo (**Fig. 4F**).

### 3.5 Chloroquine-modified siRNA-L_2_ conjugates exhibit similar *in vivo* pharmacokinetics and safety profile *in vivo* compared to parent siRNA-L_2_

Finally, we characterized the *in vivo* pharmacokinetics and safety profile of chloroquine-modified siRNA-L_2_ conjugates, particularly siRNA-CQ_2-3_-L_2_, which exhibited the best performance in the preceding *in vitro* studies. First, we compared the circulation kinetics of unlabeled siRNA-L_2_ and siRNA-CQ_3_-L_2_, injected intravenously at 1 mg/kg body weight (by siRNA weight, exclusive of the tail to ensure equal siRNA dose), quantified using a peptide-nucleic acid (PNA) hybridization assay on whole blood samples. There were no significant differences in the circulation kinetics (**Fig. 5A**) or calculated half-life (0.38 hours for siRNA-L_2_ and 0.43 hours for siRNA-CQ_3_-L_2_ using a one-phase decay model).

**Figure 5:**
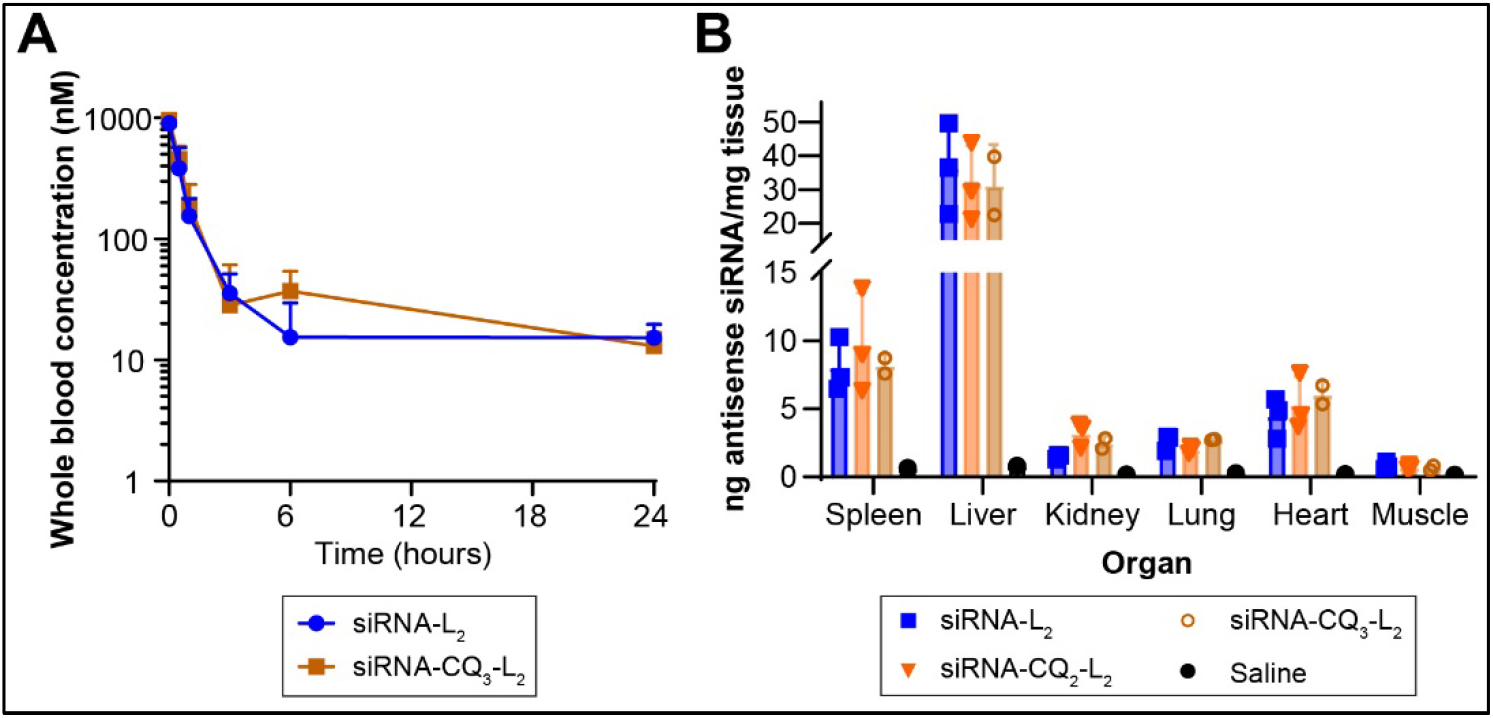
*In vivo* characterization of siRNA-CQ-L_2_ circulation kinetics and biodistribution. **(A)** Blood concentrations of siRNA-L_2_ and siRNA-CQ_3_-L_2_ after intravenous injection, quantified by PNA assay. **(B)** Organ biodistribution of siRNA-L_2_ vs. siRNA-CQ_2_-L_2_ or siRNA-CQ_3_-L_2_ vs. saline at 48 hours, quantified by PNA assay.

To establish the biodistribution and safety profile of siRNA-CQ-L_2_, we assessed complete blood counts, serum chemistries, organ pathology (by H&E stain), and siRNA accumulation (by PNA assay) 48 hours after an intravenous dose of 5 mg/kg of siRNA-CQ_2_-L_2_ or siRNA-CQ_3_-L_2_, previously established to be an effective dose in models of triple negative breast cancer, compared to siRNA-L_2_ or saline injection control.^10^ There were no noted abnormalities in blood counts or serum renal or hepatic markers (**Fig. S5**). Finally, distribution into healthy organs was essentially unchanged compared to parent siRNA-L_2_, with the highest accumulation being in the liver and spleen, with no statistically significant differences noted (**Fig. 5B**), and H&E histology of key organs did not reveal any signs of toxicity (**Fig. S6**).

## 4. Discussion

In this work, we have demonstrated the successful synthesis of a chloroquine phosphoramidite and subsequent incorporation into the albumin-binding siRNA-L_2_ construct to generate a direct siRNA conjugate containing a dedicated endosomolytic domain with tunable valency. Placement of the CQ groups between the siRNA strand and the branching tail structure, intentionally keeping the optimized ethylene glycol spacers intact, preserved the albumin-binding property of the constructs. The K_D_ binding constants were similar for siRNA-CQ_n_-L_2_ regardless of the number of chloroquine molecules, though there was a slight increase in the dissociation constant that is unlikely to have practical consequence, especially given the near-identical circulation kinetics and organ biodistribution profile compared to siRNA-L_2_ as tested in mouse models. The small difference in albumin dissociation behavior could possibly be related to the influence of chloroquine’s intrinsic weak interaction with albumin, though further studies would be needed to confirm this effect. While addition of the chloroquine makes the siRNA-L_2_ tail more hydrophobic, the overall siRNA duplex remains hydrophilic, and we did not observe any substantial change in the critical micelle concentration of the conjugates bearing CQ.

Addition of at least 2 chloroquine molecules on each strand was sufficient to produce a statistically significant improvement in endosomal disruption compared to the parent siRNA-L_2_ conjugate as measured using the Gal8 assay. While the Gal8 foci produced were generally similar in appearance to those seen with a commercial cationic lipid transfection reagent (lipofectamine), we noted a different timecourse of appearance of these foci, with lipofectamine causing endosomal disruption within 2 hours while siRNA-CQ-L_2_ produced signal more gradually over 24 hours. This is consistent with the known function of chloroquine as an autophagy inhibitor, preventing lysosomal maturation and fusion, which is a distinct mechanism of action compared to cationic lipids or polymers. Given concerns about excessive endosomal bursting leading to non-specific toxicity, inflammation, and off-target mRNA modulation,^8,18^ this contrasting endosomal escape behavior may be favorable. These same formulations with at least two chloroquine molecules also resulted in greater cell uptake *in vitro*, which was not solely attributable to greater non-specific binding to cells as evidenced by the degree of cell association when treated at 4 °C. The contrasting effect on cellular uptake seen with addition of free CQ compared to conjugated CQ, wherein the former led only to transient (∼1 hour) improvement in siRNA-L_2_ uptake while the latter resulted in more durable increases in uptake, suggests that this phenomenon may be in part attributable to greater retention rather than simply greater internalization. Corresponding with the greater uptake, we observed a trend towards greater knockdown of reporter luciferase with increasing CQ content, with siRNA-CQ_3_-L_2_ in particular producing statistically significantly greater knockdown compared to parent siRNA-L_2_ at every dose level tested. In those conditions, we measured ∼60% silencing at the protein level after just 24 hours of treatment at 1000 nM, levels of silencing that siRNA-L_2_ typically achieves at 48-72 hours when delivered “forward” over adherent cells, consistent with an acceleration of cytoplasmic delivery with the addition of sufficient quantities of chloroquine. While relatively high doses were still required to achieve meaningful levels of silencing as compared to typical doses used with traditional transfection reagents, siRNA-CQ_n_-L_2_ was very well-tolerated and did not cause any change in cell viability at the doses and timepoints tested.

In mouse models, we saw similar circulation kinetics with the siRNA-CQ-L_2_ conjugates compared to the parent siRNA-L_2_, consistent with intact albumin binding. Of note, we measured circulation kinetics using a peptide-nucleic acid hybridization assay, which avoids potential confounding effects of adding fluorescent dyes for tracking the siRNA. In contrast to prior pharmacokinetic measurements for siRNA-L_2_ made using intravital microscopy,^9,10^ we were able to establish absolute concentrations of unlabeled siRNA in the blood, with the t_0_ values of ∼1 μM consistent with the 1 mg/kg (∼2 nmol for a 25 g mouse) dose in a circulating blood volume of ∼8% body weight (2 mL for a 25 g mouse); this accuracy may permit more accurate future modeling of exposure of tumors to the siRNA conjugate therapeutics. We saw no significant differences in healthy organ biodistribution of siRNA-CQ_2_-L_2_ or siRNA-CQ_3_-L_2_ compared to siRNA-L_2_. The propensity for accumulation in the liver and spleen has been previously established and is consistent with reticuloendothelial system clearance of the siRNA-albumin complexes, in contrast with renal clearance of free siRNA.^10^ More importantly, we did not see a shift in organ distribution as a result of the positive charges added by the chloroquine molecules, which has been described in the context of lipid nanoparticles.^19^ Chloroquine bears two amines with pKa values (8.4 and 10.2) above the pH of blood and thus will on average carry two positive charges per molecule in physiologic conditions; however, the net charge of the conjugate and siRNA-albumin complex as a whole remains overwhelmingly negative.

The general synthetic approach devised for the synthesis of siRNA-CQ-L_2_ is highly modular. The number of endosomolytic moiety repeats can be precisely defined, as well as the position of these repeats relative to other parts of the conjugate. Further, the specific endosomolytic moiety can be easily exchanged. Aside from the straightforward single-step phosphoramidite synthesis reaction, which is generally applicable to any hydroxyl-containing compound of interest, production of these siRNA conjugates is entirely an on-column solid phase synthesis. Thus, in the future, the approach can be rapidly adapted to explore not only other endosomolytic configurations but also other functionalities such as targeting or stimulus-responsive systems.

## 5. Acknowledgements

We would like to acknowledge assistance from and use of the Vanderbilt University Mass Spectrometry Research Center, Vanderbilt Institute of Nanoscale Science and Engineering (VINSE) Analytical Core, Vanderbilt Center for Structural Biology (CSB) Biophysical Instrumentation Facility, Vanderbilt University Medical Center (VUMC) Molecular Cell Biology Resource, VUMC Digital Histology Shared Resource, and VUMC Translational Pathology Shared Resource (NCI/NIH P30 CA068485). We are thankful for grant support from the NIH (F32 CA268705, J.H.L.; T32 CA217834, J.H.L.; K12 CA090625, J.H.L.; R01 CA260958: M.J.U. and C.L.D.; and GI SPORE P50 CA236733), the Phi Beta Psi Sorority Trust national research awards AWD00000652 (M.J.U.) and AWD00001248 (M.J.U.); and a Vanderbilt Institute for Clinical and Translational Research (VICTR) award VR54033.1 (M.J.U.).

## Notes

### Competing Interest Statement

The authors have declared no competing interest.

## References

1. Pecot CV, Calin GA, Coleman RL, Lopez-Berestein G, Sood AK. RNA interference in the clinic: challenges and future directions. Nat Rev Cancer 2011;11(1):59–67. (10.1038/nrc2966).

2. Jackson MA, Patel SS, Yu F, et al. Kupffer cell release of platelet activating factor drives dose limiting toxicities of nucleic acid nanocarriers. Biomaterials 2021;268:120528. DOI: 10.1016/j.biomaterials.2020.120528.

3. Weiss AM, Lopez MA, 2nd, Rawe BW, et al. Understanding How Cationic Polymers’ Properties Inform Toxic or Immunogenic Responses via Parametric Analysis. Macromolecules 2023;56(18):7286–7299. DOI: 10.1021/acs.macromol.3c01223.

4. Allerson CR, Sioufi N, Jarres R, et al. Fully 2’-modified oligonucleotide duplexes with improved in vitro potency and stability compared to unmodified small interfering RNA. J Med Chem 2005;48(4):901–4. DOI: 10.1021/jm049167j.

5. Sehgal I, Eells K, Hudson I. A Comparison of Currently Approved Small Interfering RNA (siRNA) Medications to Alternative Treatments by Costs, Indications, and Medicaid Coverage. Pharmacy (Basel) 2024;12(2). DOI: 10.3390/pharmacy12020058.

6. Brown CR, Gupta S, Qin J, et al. Investigating the pharmacodynamic durability of GalNAc-siRNA conjugates. Nucleic Acids Res 2020;48(21):11827–11844. DOI: 10.1093/nar/gkaa670.

7. Bartlett DW, Davis ME. Insights into the kinetics of siRNA-mediated gene silencing from live-cell and live-animal bioluminescent imaging. Nucleic Acids Res 2006;34(1):322–33. DOI: 10.1093/nar/gkj439.

8. Biscans A, Ly S, McHugh N, Cooper DA, Khvorova A. Engineered ionizable lipid siRNA conjugates enhance endosomal escape but induce toxicity in vivo. Journal of controlled release : official journal of the Controlled Release Society 2022;349:831–843. DOI: 10.1016/j.jconrel.2022.07.041.

9. Sarett SM, Werfel TA, Lee L, et al. Lipophilic siRNA targets albumin in situ and promotes bioavailability, tumor penetration, and carrier-free gene silencing. Proc Natl Acad Sci U S A 2017;114(32):E6490–E6497. DOI: 10.1073/pnas.1621240114.

10. Hoogenboezem EN, Patel SS, Lo JH, et al. Structural optimization of siRNA conjugates for albumin binding achieves effective MCL1-directed cancer therapy. Nature communications 2024;15(1):1581. DOI: 10.1038/s41467-024-45609-0.

11. Bus T, Traeger A, Schubert US. The great escape: how cationic polyplexes overcome the endosomal barrier. Journal of materials chemistry B 2018;6(43):6904–6918. DOI: 10.1039/c8tb00967h.

12. Du Rietz H, Hedlund H, Wilhelmson S, Nordenfelt P, Wittrup A. Imaging small molecule-induced endosomal escape of siRNA. Nature communications 2020;11(1):1809. DOI: 10.1038/s41467-020-15300-1.

13. Yu F, Xie Y, Wang Y, Peng ZH, Li J, Oupicky D. Chloroquine-Containing HPMA Copolymers as Polymeric Inhibitors of Cancer Cell Migration Mediated by the CXCR4/SDF-1 Chemokine Axis. ACS Macro Lett 2016;5(3):342–345. DOI: 10.1021/acsmacrolett.5b00857.

14. Kilchrist KV, Dimobi SC, Jackson MA, et al. Gal8 Visualization of Endosome Disruption Predicts Carrier-Mediated Biologic Drug Intracellular Bioavailability. ACS Nano 2019;13(2):1136–1152. DOI: 10.1021/acsnano.8b05482.

15. Fletcher RB, Stokes LD, Kelly IB, 3rd, et al. Nonviral In Vivo Delivery of CRISPR-Cas9 Using Protein-Agnostic, High-Loading Porous Silicon and Polymer Nanoparticles. ACS Nano 2023;17(17):16412–16431. DOI: 10.1021/acsnano.2c12261.

16. Godinho B, Gilbert JW, Haraszti RA, et al. Pharmacokinetic Profiling of Conjugated Therapeutic Oligonucleotides: A High-Throughput Method Based Upon Serial Blood Microsampling Coupled to Peptide Nucleic Acid Hybridization Assay. Nucleic acid therapeutics 2017;27(6):323–334. DOI: 10.1089/nat.2017.0690.

17. Krieg B, Hirsch M, Scholz E, et al. New Techniques to Assess In Vitro Release of siRNA from Nanoscale Polyplexes. Pharm Res 2015;32(6):1957–74. DOI: 10.1007/s11095-014-1589-7.

18. Omo-Lamai S, Wang Y, Patel MN, et al. Lipid Nanoparticle-Associated Inflammation is Triggered by Sensing of Endosomal Damage: Engineering Endosomal Escape Without Side Effects. bioRxiv 2024. DOI: 10.1101/2024.04.16.589801.

19. Cheng Q, Wei T, Farbiak L, Johnson LT, Dilliard SA, Siegwart DJ. Selective organ targeting (SORT) nanoparticles for tissue-specific mRNA delivery and CRISPR-Cas gene editing. Nat Nanotechnol 2020;15(4):313–320. DOI: 10.1038/s41565-020-0669-6.

