## Supplemental Figures for "Synthesis and characterization of chloroquine-modified albumin-binding siRNA-lipid conjugates for improved intracellular delivery and gene silencing in cancer cells"

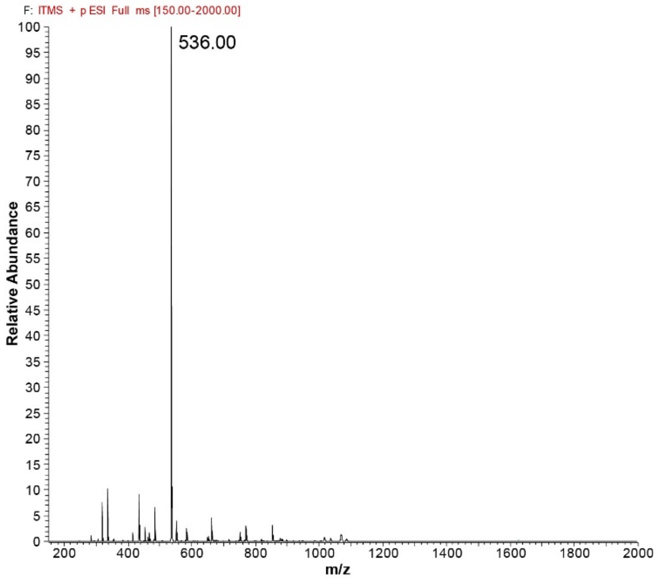


**Figure S1.** Mass spectrometry (ESI positive ion mode) confirming molecular weight of chloroquine phosphoramidite (peak shown represents [M+H])

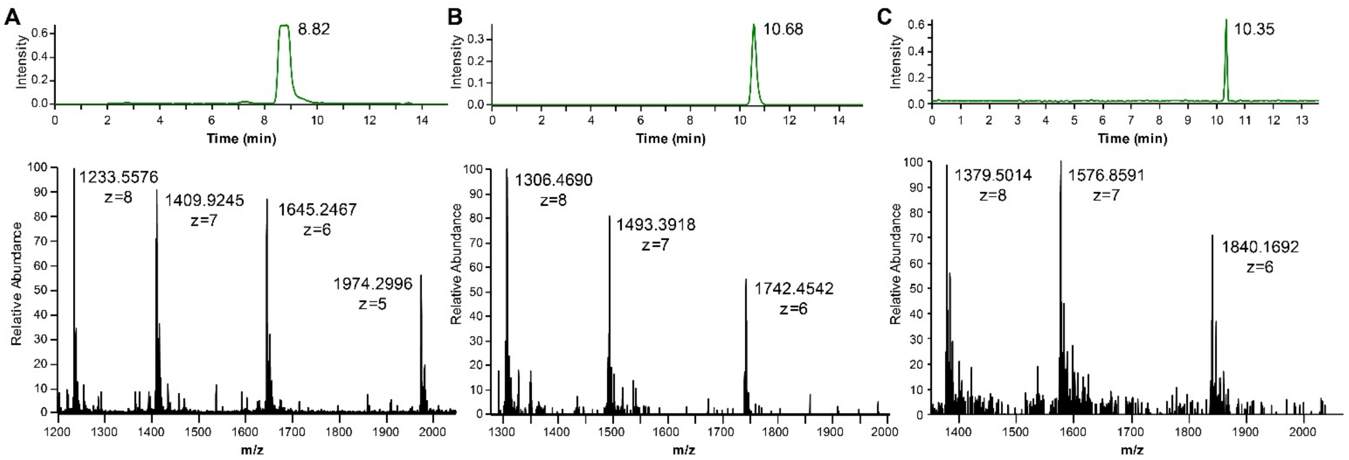


**Figure S2.** Liquid chromatography-mass spectrometry (LC-MS) confirmation of purity and molecular weight of (A) siRNA-$CQ_1$-$L_2$, (B) siRNA-$CQ_2$-$L_2$, and (C) siRNA-$CQ_3$-$L_2$ sense strands targeting luciferase, with LC chromatograms shown above and mass spectra shown below. Predicted molecular weights were 9875, 10458, and 11040 g/mol, respectively.

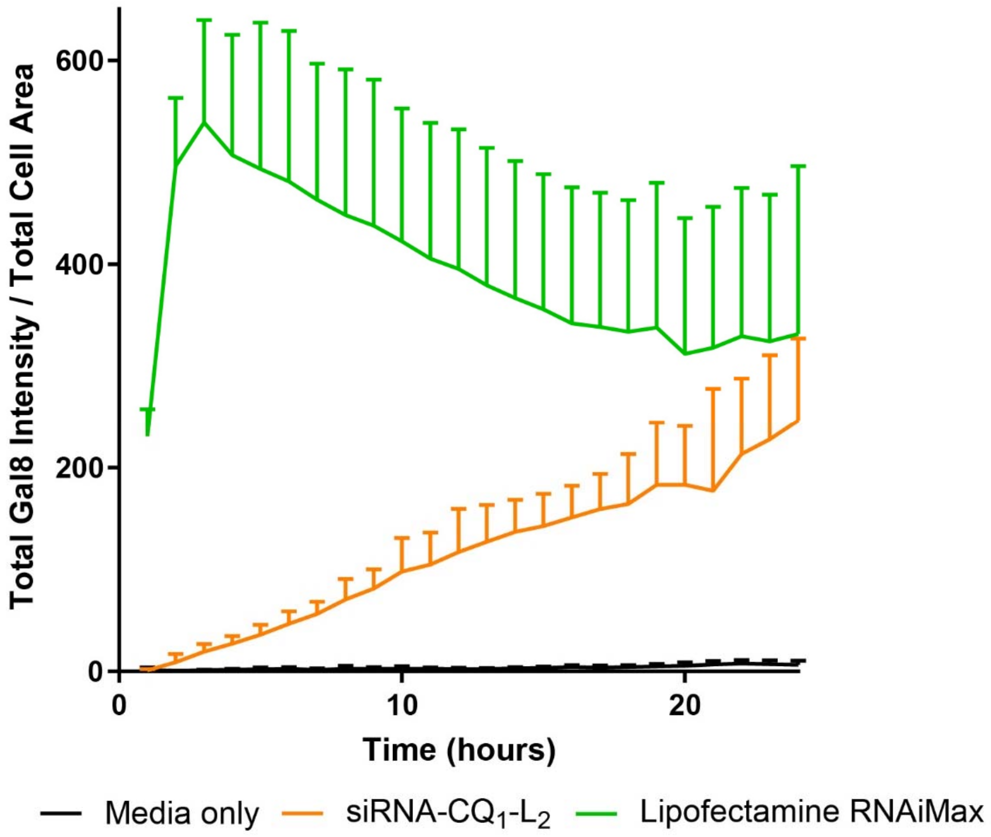


**Figure S3.** Kinetics of Gal8 focus formation in response to siRNA-$CQ_1$-$L_2$ vs. lipofectamine RNAiMax transfection reagent, assessed as total Gal8 fluorescent intensity normalized to total cytoplasmic area, over the course of 24 hours.

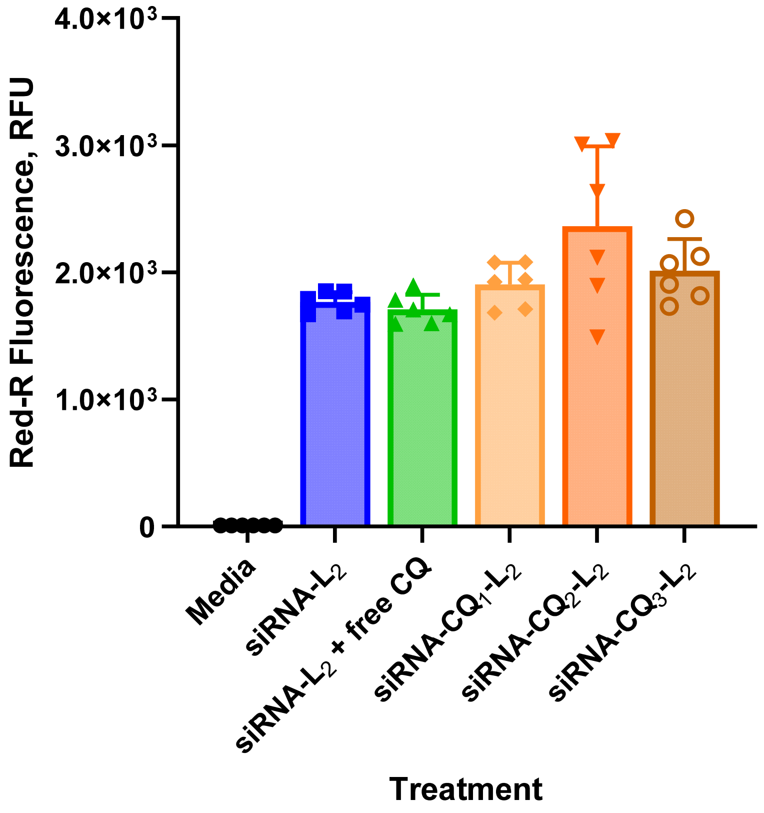


**Figure S4.** Flow cytometry analysis of uptake of Cy5-labeled siRNA-$L_2$ without or with free chloroquine or Cy5-labeled siRNA-CQ-$L_2$ in MDA-MB-231 cells incubated at 4 °C for 2 hours to assess for nonspecific binding. Geometric means of Cy5 fluorescence (n=4 per condition) are shown.

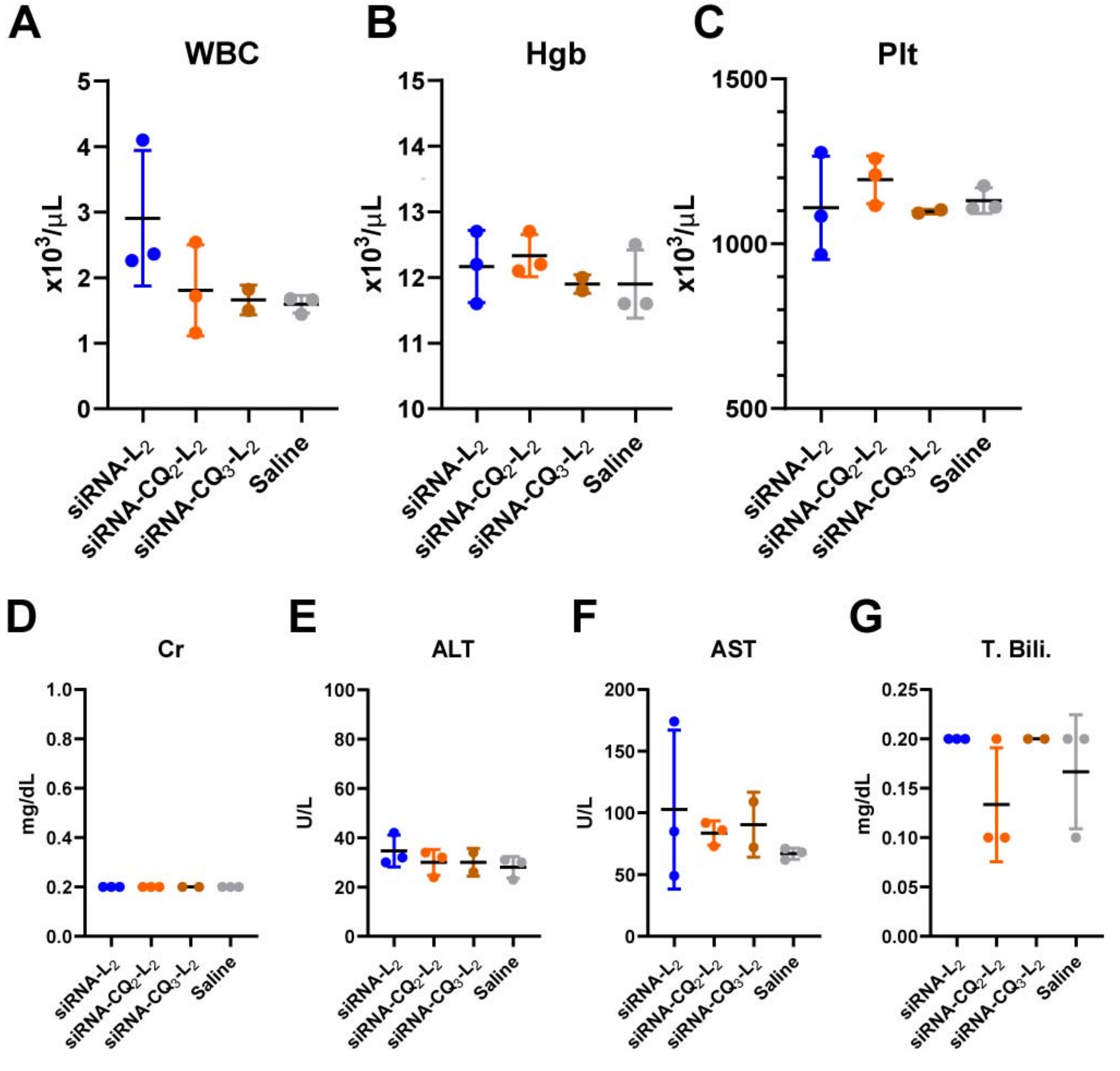


**Figure S5. (A)** White blood cell counts, **(B)** hemoglobin, **(C)** platelet counts, **(D)** creatinine (note: all values were reported as <0.2 mg/dL), **(E)** alanine aminotransferase, **(F)** aspartate aminotransferase, and **(G)** total bilirubin 48 hours following i.v. injection with siRNA-L$_2$ vs. siRNA-CQ$_2$-L$_2$ or siRNA-CQ$_3$-L$_2$ dosed at 5 mg siRNA/kg body weight.

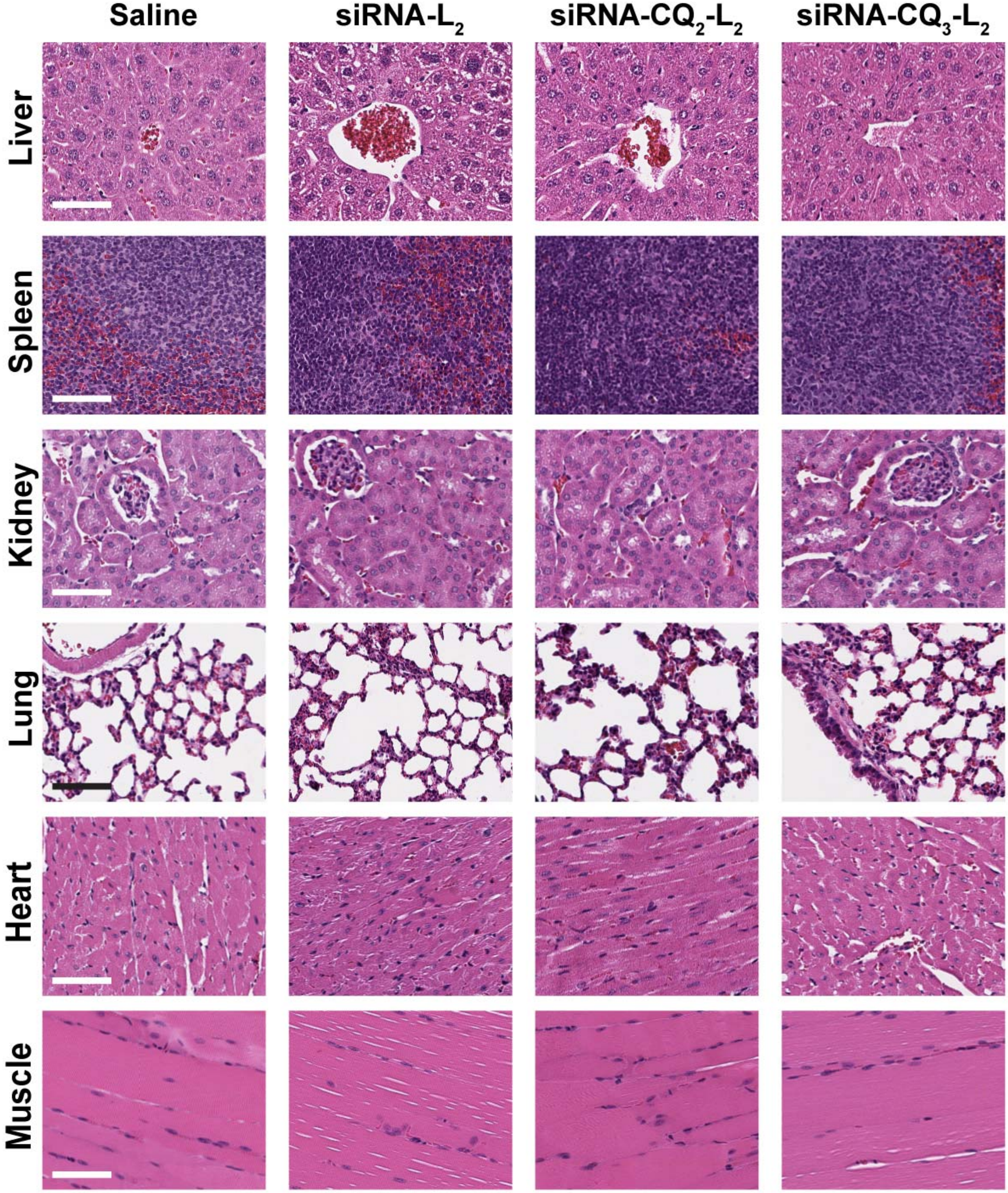


**Figure S6.** H&E histology 48 hours following i.v. injection with siRNA-$L_2$ vs. siRNA-$CQ_2$-$L_2$ or siRNA-$CQ_3$-$L_2$ dosed at 5 mg siRNA/kg body weight. Scale bar = 100 µm.
